# Low marine food levels mitigate high migration costs in anadromous populations

**DOI:** 10.1101/577353

**Authors:** P. Catalina Chaparro-Pedraza, André M. de Roos

## Abstract

Migratory fish populations, like salmon, have dramatically declined for decades. Because of their extensive and energetically costly breeding travel anadromous fish are sensitive to a variety of environmental threats, in particular infrastructure building in freshwater streams and food declines in the ocean. Here, we analyze the effects of these two threats combined. Unexpectedly, we find that low marine food availabilities favor, as opposed to threaten, the ecological success of endangered populations. This counterintuitive effect results from an aspect of individual energetics that individuals switching to higher food levels reach larger sizes with concomitant larger migration costs but have lower energy densities. Surprisingly, the decline of food levels in the ocean after the completion of dams may thus mitigate the risk of extinction of migratory fish populations. This highlights the need of a mechanistic understanding integrating individual energetics, life history, and population dynamics to accurately assess biological consequences of environmental change.

## Introduction

Migratory fish populations have shown dramatic declines over the last decades. In the North Atlantic, historical records of 24 migratory fish species reveal that they all decreased in abundance by more than 90% (Limburg & Waldman, 2009). Conservation of these populations is of particular concern because of their economic and cultural importance: although anadromous fish comprise on average less than 1% of the world fish species, their share in global fisheries trade currently exceeds 17% and continues to increase (FAO, 2016). Anadromous fish are exposed to a wide range of environmental influences in both freshwater and marine habitats, which makes them particularly vulnerable to environmental change. Multiple threats have contributed to the decline of anadromous populations (Limburg & Waldman, 2009), including infrastructure building in freshwater streams and food declines in the ocean. In fact, collapse of Atlantic salmon (*Salmo salar*) populations throughout North-Western Europe in pre-modern times coincided with the spread of watermills in river basins (Lenders et al., 2016). Likewise, a multidecadal decline of food abundance in the ocean occurs concurrently with the reduction of Atlantic salmon abundance (Friedland et al., 2009). While the effects of these threats separately are well documented, the cumulative and interactive impacts of them are poorly understood.

On their own, both food reductions in the ocean and infrastructure building in freshwater streams have negative impacts on fish life history by affecting individual energetics. Low food abundance in the ocean limits body growth rate during the oceanic stage resulting on average in individuals with smaller body sizes (Friedland et al., 2009) and concomitant lower fecundity (Thorpe, Miles, & Keay, 1984). On the other hand, dams and other anthropogenic structures increase the energetic costs of the breeding migration (Caudill et al., 2007; Mesa & Magie, 2006). In wild salmonid populations, these costs depend on the upriver distance and elevation traveled. Because individuals cease feeding when leaving the ocean and use only their energy reserves to meet the energetically costly upriver travel (N. Jonsson, Jonsson, & Hansen, 1997), this results in a depletion of energy reserves available for reproduction (Crossin et al., 2004). This depletion, however, varies between individuals of different body sizes: since transporting a large body upstream is relatively more energy demanding, large individuals spend disproportionally more energy than small ones (Bowerman, Pinson-Dumm, Peery, & Caudill, 2017; Glebe & Leggett, 1981; N. Jonsson et al., 1997) (See supporting information, Fig. S2). The presence of infrastructure in the freshwater habitat thus causes a disproportionate fecundity reduction of large-sized individuals in particular.

Anadromous fish species, as other migratory species, are exposed to a broad range of environmental influences owing to the use of multiple habitats during different parts of their life cycle. Because the freshwater habitat and the ocean differ in a wide variety of ecological and, particularly, nutritional conditions wild salmon individuals experience energetic changes (Gagliano & McCormick, 2007; MacFarlane, 2010; Werner & Gilliam, 1984) during the habitat switch with consequences for individual growth and fecundity. During their early life stages in freshwater, salmon individuals experience strong density–dependence and intense competition for food, while density–dependence in the ocean is subsequently more relaxed and food more abundant (N. Jonsson, Jonsson, & Hansen, 1998). The increase in food availability during the switch from freshwater to seawater leads to an increase in growth rates but a decrease in energy density in wild salmon (MacFarlane, 2010) with individuals after the habitat switch apparently allocating more energy to somatic growth and less to energy storage (Johansen, Ekli, Stangnes, & Jobling, 2001) (Fig. 1A). Such bias toward increased growth at the expense of energy storage following an increase in food also occurs in various other vertebrate (Auer, Arendt, Chandramouli, & Reznick, 2010; Sinervo & Doughty, 1996; Taborsky, 2006) and invertebrate (Kleinteich, Wilder, & Schneider, 2015; Zeller & Koella, 2016) species. Individuals of these species compensate their growth after an experimental period of food rationing and reach similar sizes as when experiencing continuously high food levels, at the expense of lower fecundity (Fig. 1B; see supporting information Table S1). Martin et al. (Martin, Heintz, Danner, & Nisbet, 2017) offered a mechanistic explanation for the increase in somatic growth at the expense of energy density after a transition from low to high food levels based on a well-established theory of individual energetics (S. a. L. M. Kooijman, 2010) (see supporting information, Energy allocation effects of the habitat switch explained by dynamic energy budget theory). Yet, it is not known how these energetic changes experienced by individuals during the habitat switch interact with the negative effects that low marine food levels and infrastructure building in freshwater streams independently have on individual energetics.

**Fig. 1.**
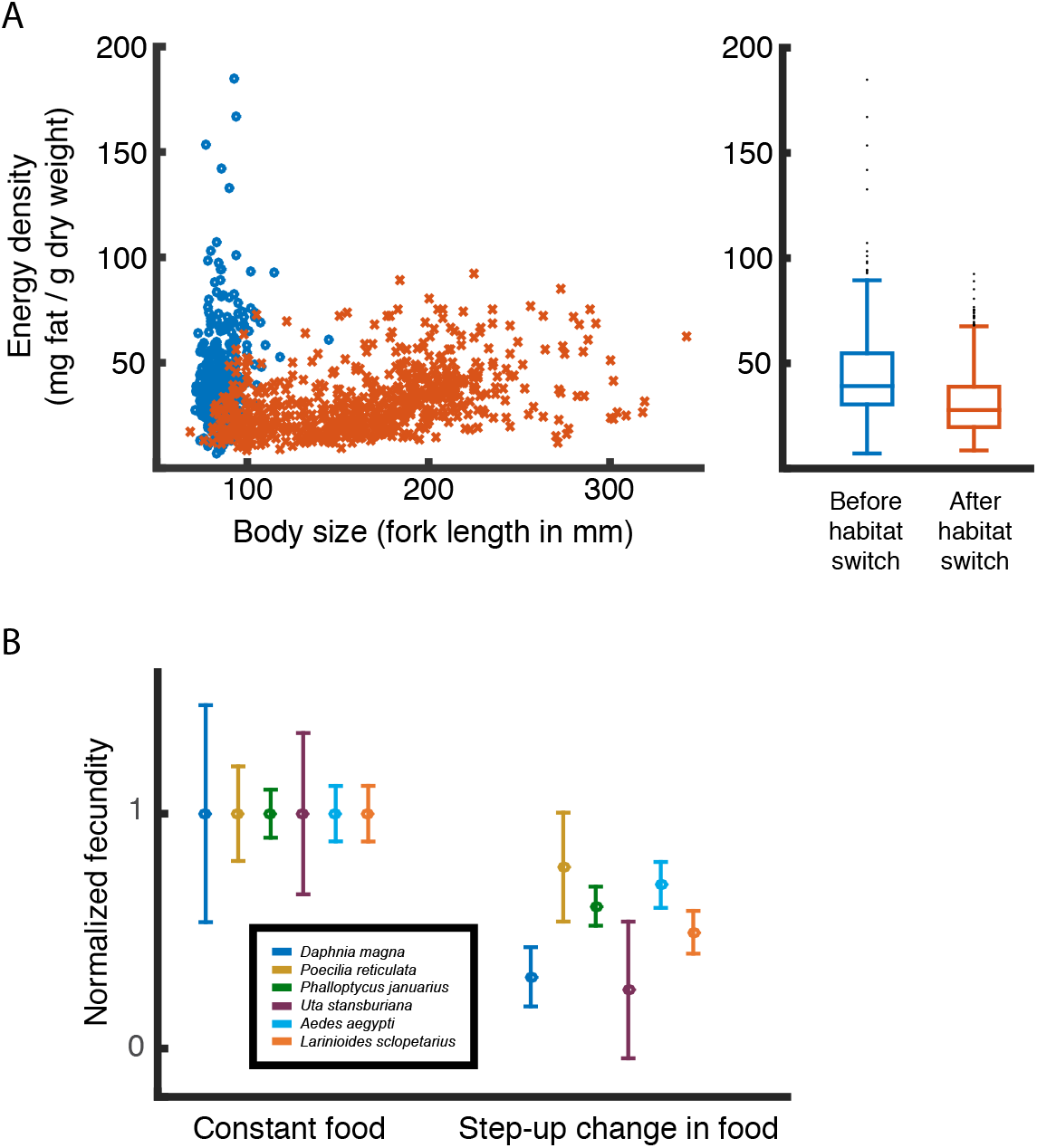
Effect of step-up change in food on energy reserves and fecundity **A**: Observed energy density as a function of individual body length in Sacramento River Chinook salmon before (blue circles) and after the habitat switch (red crosses) (Kruskal-Wallis test p<0.001)(Martin et al., 2017). The right panel summarizes the observations for individuals before and after the habitat switch independent of their length. **B**: Effect of an experimental step-up change in food provisioning on fecundity per unit of biomass in six species (Auer et al., 2010; Kleinteich et al., 2015; Sinervo & Doughty, 1996; Taborsky, 2006; Zeller & Koella, 2016) (data for *D. magna* from Kooijman, unpublished). To allow comparison between species the mean fecundity of control individuals (constant high food) is normalized to 1. The error bars represent the standard deviation.

Infrastructure building in spawning streams and declining food levels in the ocean are concurrently occurring, however these stressors have been studied only in isolation. The population consequences arising from the individual effects of these two threats combined have not been explored. In this study we assess the cumulative and interactive impact of high costs of the breeding migration and a decline in ocean food availability on the persistence of *Salmo salar* populations using a population dynamic model based on the individual energetics and life history.

## Methods

### The model

We formulate a physiologically structured population model that includes two habitats and an anadromous population. We assume each habitat to have a different food resource. The anadromous population is structured by age, structural and reversible mass and follows semi-discrete dynamics: continuous dynamics describe the resource consumption, growth in both structural and reversible mass and survival and a discrete map describes the pulse-wise reproduction (Persson, Leonardsson, de Roos, Gyllenberg, & Christensen, 1998). Structural mass we consider to include bones and organs which cannot be starved away to cover energetic demands, whereas reversible mass comprises stored energy reserves such as fat and gonads. The core part of the model is the description of the individual life history and energetics, that is, feeding, growth, reproduction and mortality as a function of the individual state (age, structural and reversible body mass) and the state of the environment (food abundance and temperature). We use data from Atlantic salmon to parameterize the model (see supporting information Table S2).

### Yearly cycle and life history events

The first life stages of Atlantic salmon occur in the freshwater habitat where the parents breed. Female salmon spawn in autumn at time *t_r_*, the eggs develop throughout winter and hatch at age *a_h_* (in spring) (Hendry & Cragg-Hine, 2003). After hatching, individuals remain in the stream until age *a_s_*, when they smolt and migrate to the sea (Hendry & Cragg-Hine, 2003) (hereafter, individuals younger than *a_s_* are referred to as presmolts, in contrast to older individuals that are referred to as postsmolts). Every autumn, all sexually mature individuals in the ocean return to the freshwater habitat and start migrating upstream at time *t_um_* to spawn somewhat later at time *t_r_*. After spawning, postsmolts return to the sea and finish downstream migration at time *t_dm_* of the year.

### Habitats

Food density in the freshwater habitat grows continuously and declines through the foraging of individuals, whereas the impact of foraging on the resource in the ocean is assumed negligible, such that food availability in this habitat is constant.

Density-dependence is strong in the breeding habitat (N. Jonsson et al., 1998) and directly affects growth in body size in salmonids (Walters, Copeland, & Venditti, 2013). We therefore incorporate density-dependent effects on the growth rate of presmolts via competition for food. In the absence of consumers, the resource is assumed to follow a semi-chemostat growth dynamics with maximum density *R_max_* and growth rate *ρ* (for an explanation and justification of this type of growth dynamics, see Persson et al. (Persson et al., 1998)). Dynamics of the resource density *R_r_* in the breeding habitat is hence given by:

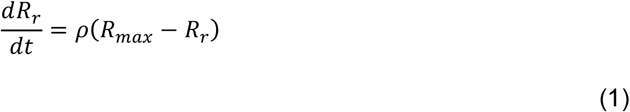

In contrast, in the marine habitat, we assume a constant feeding rate as density-dependence is negligible (N. Jonsson et al., 1998).

In both habitats, temperature *T* oscillates during the year around the average temperature *T_m_* with an amplitude *T_a_* and a period equal to the length of the year. Therefore, the temperature reaches its maximum in summer (approximate after 90 days of egg hatching) and its minimum in winter (approximate 273 days after egg hatching).

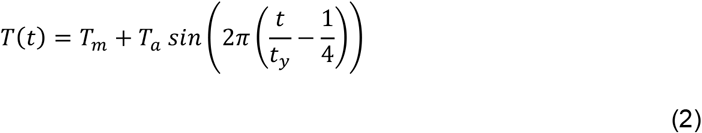

where *t* is the current time in days and *t_y_* is the number of days in a year.

### Individual dynamics

#### 1) Feeding

In the breeding habitat, individuals are assumed to feed on the resource following a Holling type II functional response. Their feeding level *f_r_* (or scaled functional response, which is the actual feeding rate at a certain food level divided by the maximum feeding rate for its current size) is given by:

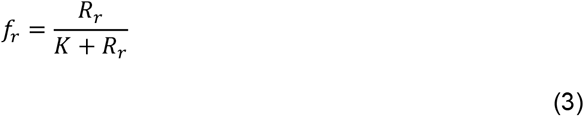

where *K* is the half saturation resource density.

In the non-breeding habitat density-dependence is assumed to be negligible so individuals feed at a constant feeding level *f_s_*.

#### 2) Dynamic energy budget model: Individual states

The description of the individual energetic dynamics and its link with life history follows the bioenergetics approach introduced by Kooijman and Metz (S. A. L. M. Kooijman & Metz, 1984) in which the energy allocation to somatic and reproductive metabolism are proportional to a fraction *κ* and a 1−*κ* of the total energy assimilation rate, respectively. Specifically, we adopt the model developed and described in detail by Martin et al. (Martin et al., 2017) for fishes; because it provides an explicit dynamics of energy reserves (in our model represented by reversible mass), which are necessary to model individual life history with starvation periods like those happening to salmon during the breeding migration. Here we provide only a concise synopsis of the model.

Individuals are characterized by three state variables: individual age *A*, structural mass *W* and reversible mass *S*. The body mass of an individual is the sum of its structural and reversible mass. Body length *l* and structural mass *W* are related to each other following:

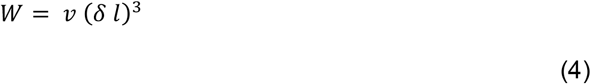

where *v* is the density of structural mass and *δ* is a shape coefficient factor. Hereafter we refer to *l* as body size (notice that body size is different to body mass described above because it only relates to structural mass).

The acquisition and utilization of energy are given by equations (5) to (13).

The energy assimilation flux is given by:

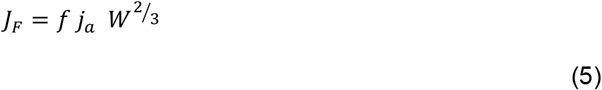

where *f* is the feeding level in either the breeding or the non-breeding habitat, *j_a_* is the maximum area-specific assimilation rate and the surface area for assimilation is assumed to scale with the structural mass to the power of 2/3.

Metabolic maintenance costs are the product of the structural mass-specific maintenance costs *j_m_* and the structural mass:

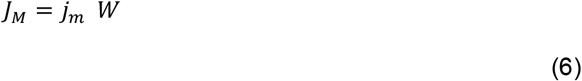

Assimilates are assumed to split into two fluxes: the *κ* flux and the 1 − *κ* flux. The *κ* flux is used to meet metabolic maintenance requirements first, while the remaining flux *J_W_* is used to synthesize structural mass. In turn, the 1 − *κ* flux *J_S_* is allocated to reversible mass. Thus,

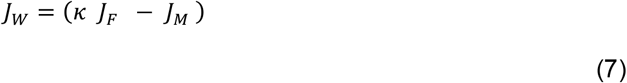

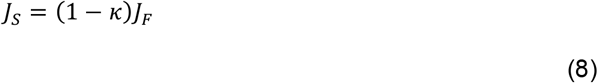

If the metabolic maintenance costs are larger than the *κ* flux, the individual starves, stops growing and thus depletes its reversible mass to cover the deficit in maintenance requirements:

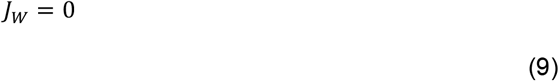

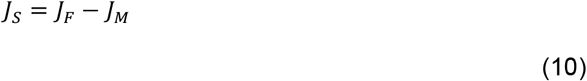

Therefore, the individual state dynamics are described by the following system of differential equations:

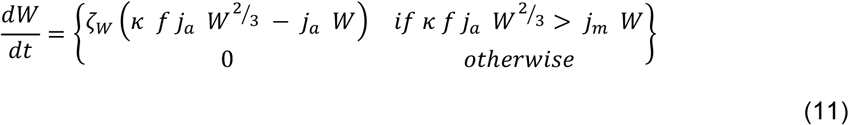

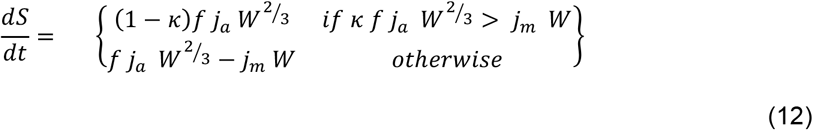

The parameter *ζ_W_* in equation (11) represents the efficiency with which assimilates are converted into structural mass.

The rate constants (*j_a_, j_m_*) are assumed to be temperature-dependent and scale from the reference temperature *T*\* to the actual temperature at time *t, T*(*t*), following the Arrhenius relationship. Hence, both rate constants are multiplied by the temperature-correction factor *F_T_*(*t*):

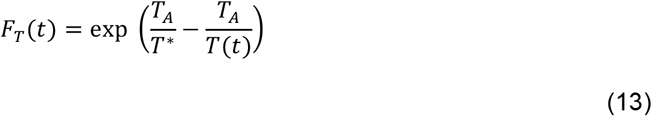

where *T_A_* is the Arrhenius temperature.

#### 3) Maturation, reproduction and breeding migration

Individuals mature when they have reached a fixed structural mass *W_p_*. The reversible:structural mass ratio at maturity *S_p_*/*W_p_* is the threshold for reproductive investment; therefore, the surplus of reversible mass in excess of the amount that equals the *S_p_*/*W_p_* reversible:structural mass ratio is used for reproduction. Reproduction occurs at a discrete time *t_r_*. The number of offspring *θ* produced by an adult individual with structural mass *W* and reversible mass *S*, hence, equals:

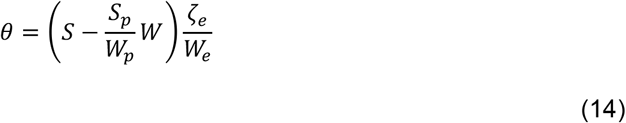

Simultaneously, the reversible mass of reproducing individuals is reduced to *S* = (*S_p_*/*W_p_*) *W*. The number of offspring produced is dependent on the yield for the conversion of reversible mass into eggs *ζ_e_* and the egg mass *W_e_*. During the egg stage individuals do not feed, therefore we assumed newly hatched individuals to be born at *t_h_* with a structural mass equal to *κ W_b_* and a reversible mass (1 − *κ*) *W_b_*.

During the breeding migration the individuals cease feeding, stop growing and use their reversible mass to meet their metabolic maintenance costs and the costs of the travel (see further justification of breeding migration costs in supporting information, Size-scaling costs of the breeding migration with structural mass), which is assumed to be proportional to the metabolic maintenance:

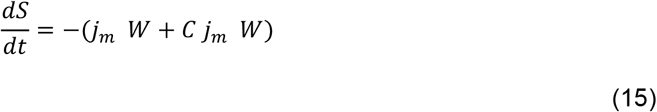

where *C* represents the relative costs of the breeding migration. According to equation (15) structural mass-specific energetic costs of the breeding migration are the same for every individual regardless of their body size, while total body mass-specific energetic costs of the breeding migration decrease with body size as reversible mass increases with body size (supporting information, Fig. S4).

#### 4) Survival

Individuals may die from either starvation or background mortality. Starving individuals with an *S*/*W* ratio smaller than *q_s_* die at a rate:

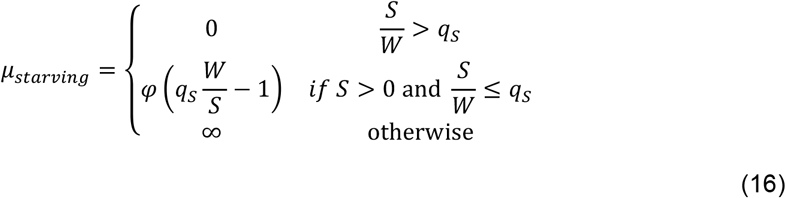

where φ is a positive proportionality constant (Persson et al., 1998).

Once they have depleted their reversible mass entirely (*S* = 0), starving individuals die instantaneously.

In addition to starvation mortality, individuals die at a rate *μ_e_* during the egg stage, at a rate *μ_r_* if they are presmolts and *μ_s_* if they are postsmolts. The total per capita death rate is the sum of the different sources of mortality.

### Population dynamics

The migratory population is structured by age, structural and reversible body mass following the approach of Persson et al. (Persson et al., 1998). Since reproduction occurs as a discrete event at a specific time in the year, all individuals that are born in the same reproductive event are lumped into a single cohort and assumed to grow at the same rate. Hence, at the population level, we can describe the dynamics of every cohort by using a system of ordinary differential equations, which keeps track of the number of individuals, their age, structural and reversible mass (see supporting information, Physiologically–structured population model). Therefore, the dynamics of the population can be followed by numerically integrating the ordinary differential equations for each cohort separately. When a reproductive event occurs, a new cohort is added to the population, which implies additional differential equations describing the population dynamics. In addition, changes in food density in the freshwater habitat can be followed by numerical integration of the ordinary differential equation that accounts for resource food growth and consumption. The numerical integration is carried out using the Escalator Boxcar Train (de Roos, 1988), a numerical integration method specifically designed to handle the system of differential equations that describes a physiologically structured population model.

### Model analysis

In the model, we investigate the effect of a decline in food availability in the ocean under both scenarios of high and low costs of the breeding migration. To do so, we computed the dynamics of the population biomass for 70 years exposed to low (*C* = 0.5) and high costs of the breeding migration (*C* = 1.5). During the first 20 years the feeding level in the ocean *f_s_* was assumed high (0.9 times the amount of food *ad libitum*) whereas it was low during the following 50 years (0.5 times the amount of food *ad libitum*) (Fig. 2A). In this simulation, the survival probability of postsmolts *μ_s_* was equal to 0.1. In addition, we studied dynamics for feeding levels in the ocean *f_s_* between *ad libitum* food and 0.5 times *ad libitum* food and costs of the breeding migration *C* between 0 and 2 times the metabolic maintenance costs with survival probability of postsmolts *μ_s_* equal to 0.1 (Fig. 4A). For the same range of feeding level *f_s_* used in Fig. 4a, we investigate the effect of variation in the background mortality rate of postsmolts *μ_s_* between 0.0063 and 0.0107 per day, equivalent to annual survival probabilities of 0.1 and 0.02, respectively (Fig. 4B). In addition, we investigate the effect of a nonlinear scaling of the costs of the breeding migration with structural mass and of reversible mass contributing to the costs of the breeding migration (see supporting information, Size-scaling of the breeding migration costs with structural mass and breeding migration costs dependent on structural and reversible mass). The effect of a nonlinear scaling of the costs of the breeding migration with structural mass is investigated with a variant of equation (15):

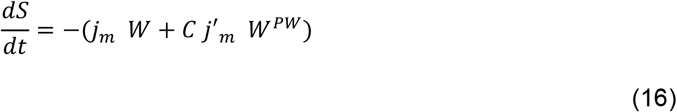

where *PW* is the size scaling exponent for the costs of the breeding migration, which in the original model had a default value equal to 1; and *j′_m_* is the structural mass-specific metabolic costs of the breeding migration. The effect of non-linear scaling of the costs of the breeding migration with structural mass is investigated for feeding levels in the ocean *f_s_* between *ad libitum* food and 0.5 times *ad libitum* food and different values of the scaling constant *C* (see supporting information, Fig. S3). The effect of reversible mass contributing to the costs of the breeding migration is investigated using the following variant of equation (15):

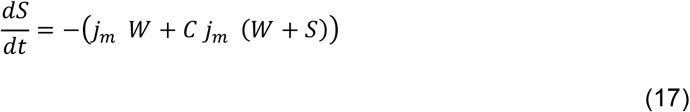

**Fig. 2.**
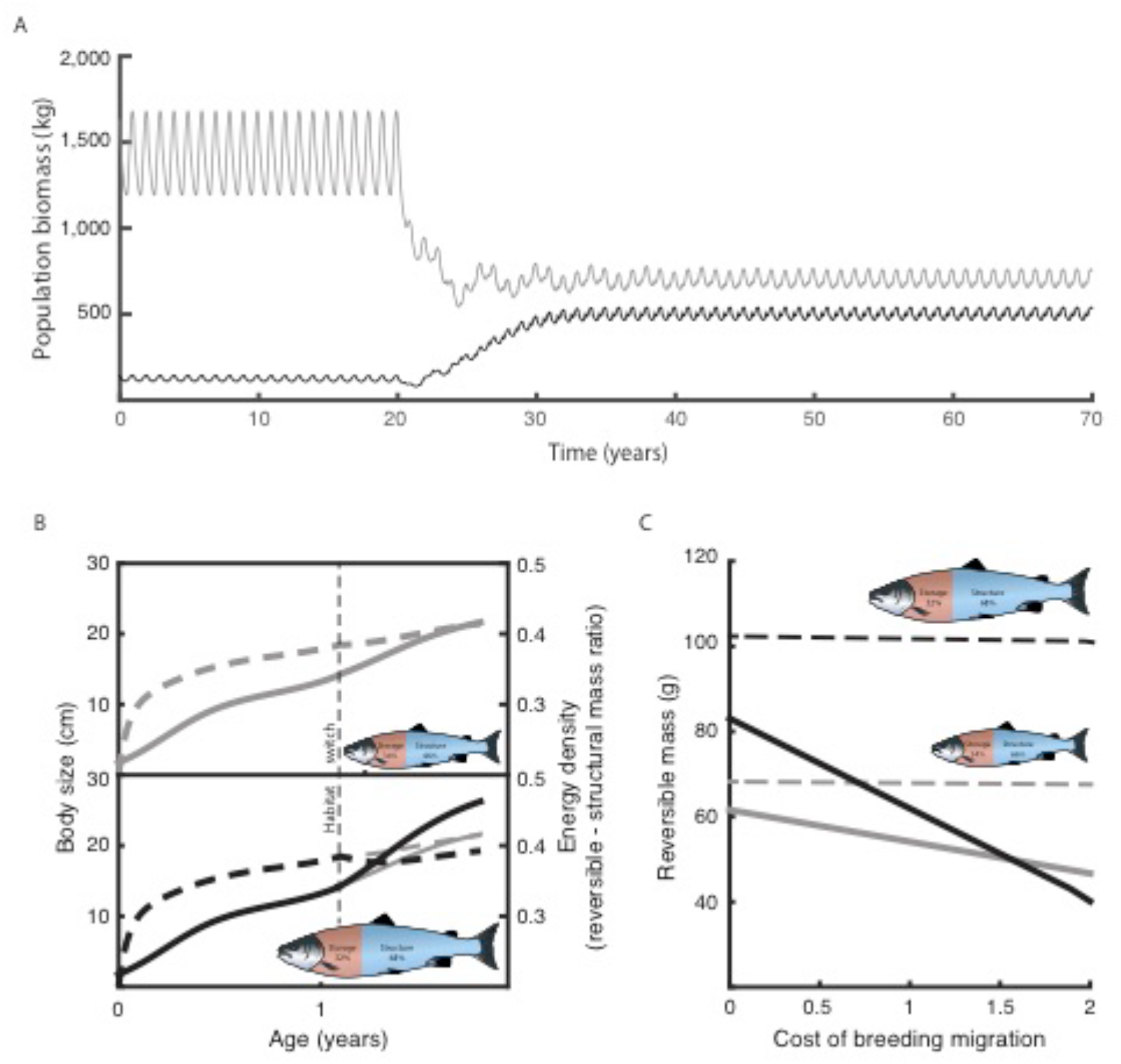
Population and individual consequences of declining food abundance in the ocean **A**: Modeled biomass dynamics of a *Salmo salar* population facing low (0.5 times the metabolic maintenance costs; grey) and high (1.5 times the metabolic maintenance; black) costs of the breeding migration preceding and following a drop in feeding level in the ocean (90% of maximum food intake during year 0-20 years and 50% of maximum food intake during year 20-70). **B**: Body size (solid line) and energy density dynamics (i.e. ratio between reversible and structural mass; dashed line) from hatching till the start of the first breeding migration for two individuals exposed to either small (feeding level changes from 50 to 60% of maximum food intake during the habitat switch; gray lines in top panels, also shown in bottom panels for comparison) or a large step-up change in food availability (feeding level change from 50 to 90% of maximum food intake during the habitat switch; black lines in lower panel). The fishes show the relative sizes and body compositions of individuals before their first breeding migration. **C**: Total reversible mass of the same individuals before (dashed line) and after (solid line) their first breeding migration dependent on the costs of the breeding migration (expressed as multiple of the metabolic maintenance costs when not travelling).

The dynamics of the population biomass is computed for 70 years exposed to low (C = 0.5) and high costs of the breeding migration (C = 1) and a decrease in the feeding level in the ocean in the year 20 as in Fig. 2 by using equations (15) and (17). The high costs of the breeding migration are lower than in Fig. 2 because when reversible mass contributes to the costs of the breeding migration and C = 1.5 the population is extinct when the feeding level in the ocean is 0.9 (see supporting information, Fig. S5).

## Results

A decline in food availability in the ocean decreases the biomass of populations when the costs of the breeding travel are low or intermediate, but surprisingly increases the population biomass when individuals experience high travelling costs (Fig. 2A) due to more extensive infrastructure in the freshwater habitat. This counterintuitive outcome results from the size-dependency of the travelling costs and the smaller energy allocation to growth in structural mass compared to reversible mass when food availability in the ocean is low. With an increase in food, growth in structural mass increases more than growth in reversible mass and this bias to structural mass growth scales with the magnitude of the food increase. Consequently, low food abundance in the ocean will result in lower structural mass and thus smaller, but fattier salmon individuals after their habitat switch (Fig. 2B; see supporting information, Energy allocation effects of the habitat switch explained by dynamic energy budget theory). Their smaller sizes entail lower energy requirements during their first travel to the spawning grounds, while relative to their body size they have more energy reserves to support them. In contrast, individuals experiencing a large increase in food availability when switching habitats are larger but have lower reversible-structural mass ratios and thus require proportionally more of their energy reserves for their breeding travel. When costs of the breeding travel are high, individuals with smaller body sizes but higher reversible-structural mass ratios suffer a less dramatic depletion of their energy reserves and thus have more energy available for reproduction upon arrival at the spawning grounds (Fig. 2C). In contrast, when the costs of the breeding travel are lower, larger individuals, having more reversible mass (in absolute value, but lower relative to their body size), arrive with more reversible mass at the spawning grounds and thus have higher fecundity. With lower costs of the breeding travel the depletion of energy reserves is not as dramatic and a decline in food availability in the ocean hence decreases population biomass. In summary, high costs of the breeding travel favor small individuals over large, and conversely, low costs favor large over small.

Our population dynamic model also predicts that the highest population biomass occurs, as expected, in unthreatened anadromous populations: when energetic costs of the breeding migration are low and food levels and survival in the ocean are high. Conversely, it predicts that populations facing high energetic costs of the breeding migration or low survival in the ocean go extinct when food availability in the ocean, and therefore the growth potential, is high (Fig. 3). However, under these stressed conditions low food abundance in the ocean enables persistence of the population. The effect that population biomass decreases with declining food availability in the ocean for low to intermediate costs of the breeding migration, whereas it increases when individuals experience high migration costs, occurs robustly as long as the breeding migration costs scale with structural mass with a power larger than 0.5 and also when reversible mass contribute to the migration costs (see supporting information, Size-scaling of the breeding migration costs with structural mass and breeding migration costs dependent on structural and reversible mass, Fig. S3, S5).

**Fig. 3.**
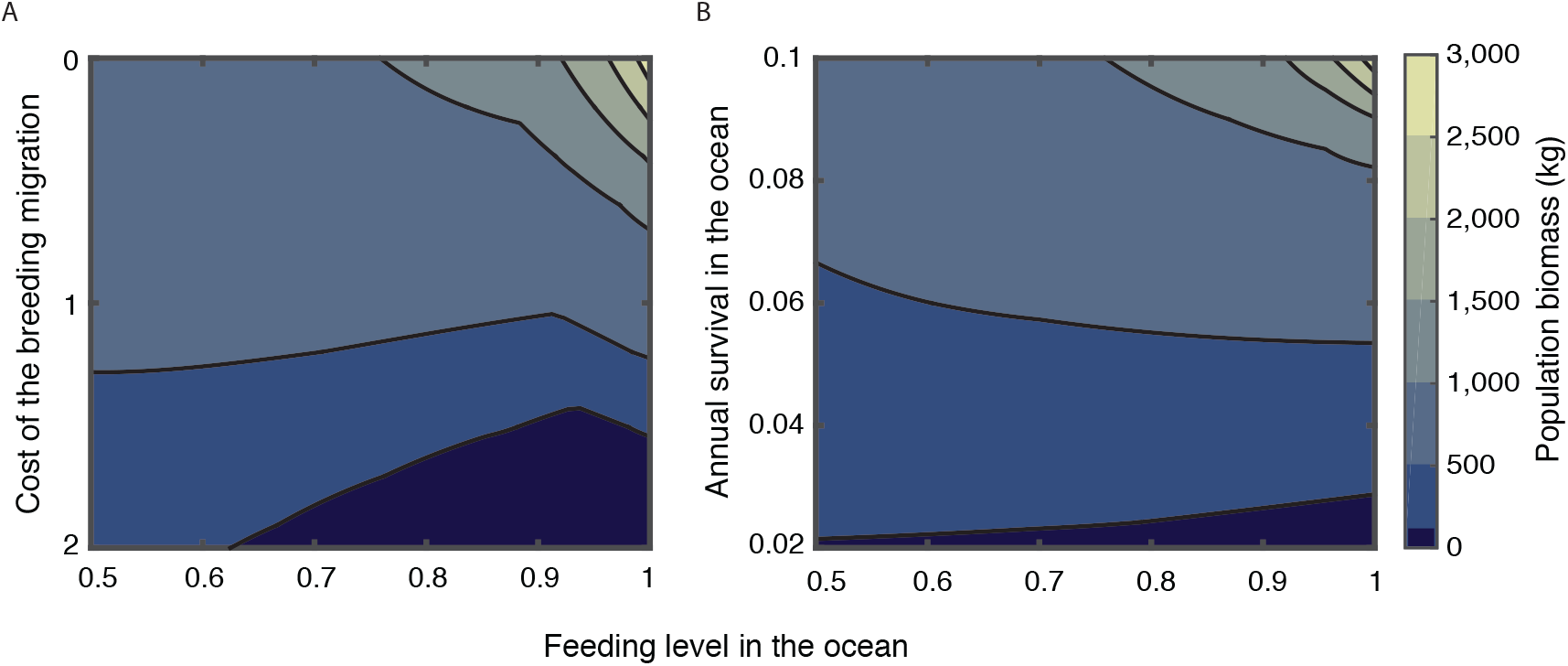
Consequences of interacting threats for population biomass Predicted biomass of anadromous populations exposed to different feeding levels in the ocean (horizontal axes) and facing different **A**: costs of the breeding migration and **B**: annual survival of migrants. A population is considered extinct when its biomass is smaller than 100 kg (dark blue). Values shown represent the average population biomass computed over the stable annual cycle that the population exhibits.

We searched for empirical observations to explore the prediction that a decline in food availability and thus in growth potential in the ocean increases the biomass of a population when costs of the breeding migration are high. In line with our results, the salmon populations from the Imsa (river length ~1 km) and the North Esk river (river length>100 km) have shown a decrease and increase in abundance, respectively, while the body growth in the ocean has decreases in the same period in both populations (Fig. 4).

**Fig. 4.**
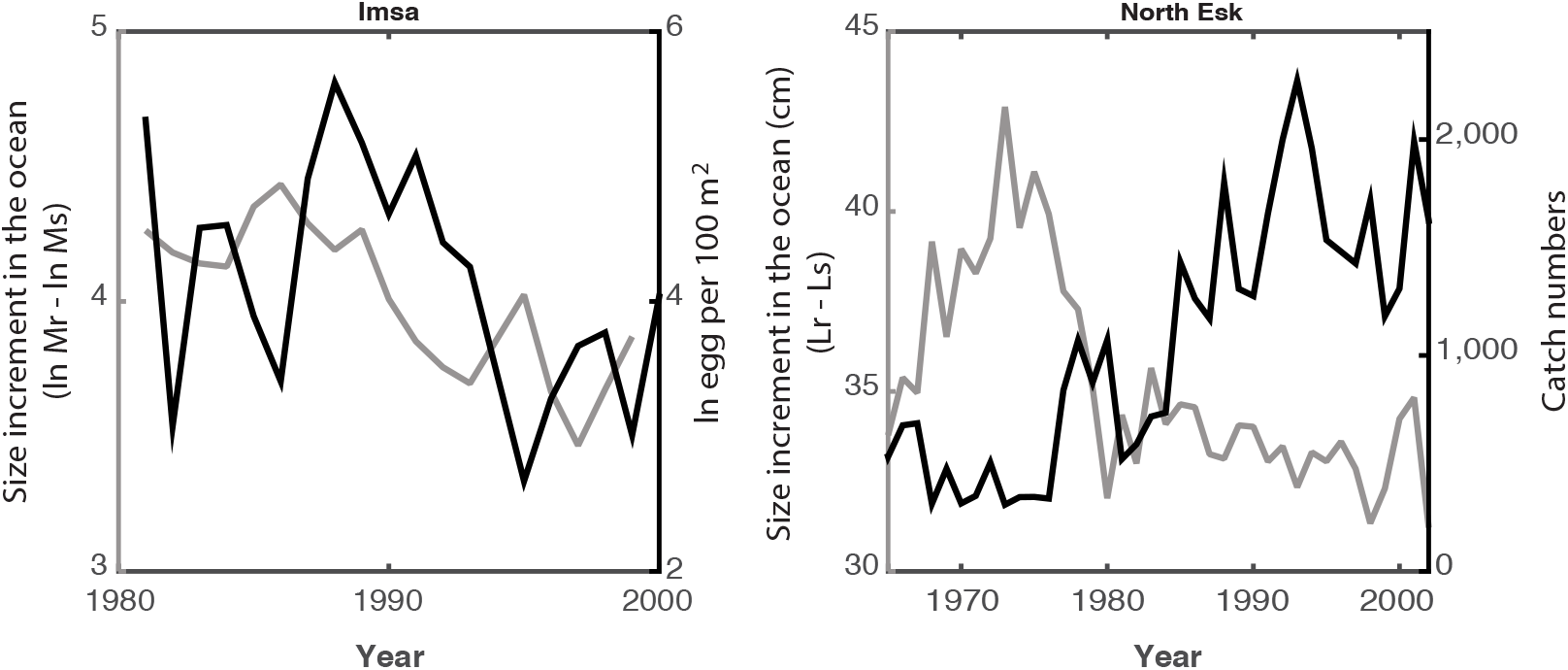
Consequences of declining food availability in the ocean in two wild Atlantic salmon populations Increase in individual body size of *Salmo salar* during its oceanic life stage (Friedland et al., 2009; B Jonsson & Jonsson, 2004) (gray) and population abundance (black) in number of eggs (Bror Jonsson & Jonsson, 2016) or catch (ASFB, 2010) from the rivers Imsa and North Esk, respectively. Mr = mass of returning adults and Ms = mass of the same fish at the habitat switch, for the same life stages, Lr and Ls correspond to the fork length.

## Discussion

The beneficial effect of low marine food availabilities on anadromous populations threatened by other stressors such as high energetic costs of the breeding migration or low survival in the ocean is caused by the highly non-linear manner in which changes experienced by individuals during the habitat switch and the effects of low marine food levels and high energetic costs of the breeding migration interact and influence individual energetics. The mechanism underlying this counterintuitive result arises from the fact that large individuals spend more energy than small ones transporting their body upstream (Bowerman et al., 2017; Glebe & Leggett, 1981; N. Jonsson et al., 1997) and that a low food level in the ocean influences the individual energetics in two ways: 1) it entails a lower growth potential and therefore a smaller body size and, 2) it causes a small increase in food availability during the habitat switch, which reduces the bias toward growth in structural versus reversible mass. As a consequence, high costs of the breeding migration favor small individuals over large, and conversely, low costs favor large over small. Any factor decreasing the individual growth rate, such as a decline in food availability, will therefore increase population biomass in the first case but decrease it in the second case. Accordingly, wild populations of sockeye salmon (*Oncorhynchus nerka*) experiencing naturally high costs of the breeding migration are composed of smaller individuals than populations facing lower costs of the breeding migration (Crossin et al., 2004).

The size-scaling of the energetic costs of the breeding migration is key in the mechanism causing the favoring effect of low marine food levels on threatened anadromous populations. In the model, we assumed the energetic costs of the breeding migration to be proportional to the structural mass of an individual and thus structural mass-specific energetic costs of the breeding migration to be the same for all individuals regardless of their body size (see supporting information, Fig. S4). Although in laminar flows the mass-specific costs of swimming are usually considered to decrease with body size (Alexander, 1998), actual field data of energy expenditure during the breeding migration of Chinook salmon *Oncorhynchus tshawytscha* show that the structural mass-specific energy requirements increase with body size (Bowerman et al., 2017). That is, larger individuals spent more energy per unit of structural mass than smaller ones and the energetic costs of the breeding migration scale hyperallometrically with structural mass (Fig. S2). Based on this evidence, our assumption of size-scaling factor is conservative, as a hyperallometric scaling of energy costs with body size would exacerbate the effect by causing large individuals to spend even more energy than small ones.

We find support for the effect that population biomass decreases with declining food availability in the ocean for low to intermediate costs of the breeding migration, whereas it increases when individuals experience high migration costs in long-term data of salmon populations from the Imsa and the North Esk rivers. In the first the population abundance has declined while in the last it has increased, despite that a declining trend in body growth in the ocean, due to low marine food availability, is observed in the populations of both rivers (Friedland et al., 2009). In accordance with our model prediction, salmon in the Imsa river indeed migrate short distances in freshwater while in the North Esk river they migrate long distances and hence face higher energetic costs of the breeding migration.

Various consequences for the conservation and fisheries industry of Atlantic salmon and other anadromous species arise from the prediction that populations facing high energetic costs of the breeding migration or low survival in the ocean go extinct when food availability in the ocean, and therefore the growth potential, is high. In particular, increased harvesting or increased costs of migration by infrastructure construction in the freshwater habitat are more likely to result in extinction of salmon populations migrating to a highly productive marine habitat, especially if these negative impacts occur in a time frame that is too short for an evolutionary response towards, for example, slow body growth rate and other physiological and morphological adaptations for an energetically costly upriver travel (Eliason et al., 2011). Although the rate at which dams were completed declined from around 1000 a year in the 1970s to around 260 a year in the 1990s, during the first decades of the 20th century the number of large dams increased with on average more than 500 new dams annually from 45000 (WCD, 2000) to 55000 (ICOLD, 2017). This trend implies that an increasing number of anadromous populations are facing higher costs of the breeding migration, possibly resulting in local extinction or selection for smaller body sizes with negative effects for fisheries in either case. Paradoxically, the decline of food levels in the Northeast Atlantic during the last decades may have mitigated the collapse of Atlantic salmon populations due to increased infrastructure building in freshwater streams. In the upcoming decades food levels are predicted to continue dropping due to climate change (Hoegh-Guldberg & Bruno, 2010; Moore et al., 2018), which although bad for fisheries, may benefit salmon population persistence.

Other factors may affect the persistence of anadromous populations facing concurrently high costs of the breeding migration and declining food availability in ocean. After the habitat switch, size-dependent predation mortality risk increases and therefore high food availability in the ocean that enables high body growth rate increases survival in the first marine stage (Friedland et al., 2009) favoring the persistence of threatened populations. Reduced size-dependent predation mortality risk could be the result either of a high growth rate during the first marine stage or of reaching a large body size before the habitat switch. In fact, a longterm study of Atlantic salmon in the Simojoki river shows that survival during the marine phase is positively correlated with body size at the habitat switch (Jutila, Jokikokko, & Julkunen, 2006). Moreover, the body size at the habitat switch in this population is negatively correlated with the individual density in the freshwater habitat (Jutila et al., 2006); therefore large body sizes at the habitat switch are reached when population abundance is low in this habitat. This is the consequence of relaxed density-dependence enabling a high body growth rate (Walters et al., 2013). Since threatened anadromous populations have relatively low population abundance, this suggests that its individuals are more likely to reach a larger body size at the habitat switch and therefore to experience lower predation mortality once in the ocean. Further research is necessary to understand how size-dependent predation mortality interacts with reduced food availability in the ocean and high costs of the breeding migration accounting for density-dependence effects in the freshwater habitat.

Current environmental changes increase the variety and intensity of stressors affecting ecological communities in cumulative and interactive ways (Crain, Kroeker, & Halpern, 2008). We demonstrate that two specific stressors with independent negative effects interact in a highly non-linear manner as a consequence of the specifics of individual energetics and life history. Given that environmental change comes with other stressors such as warming trends in the spawning streams that also affect individual salmon energetics and thus life history (Eliason et al., 2011; B. Jonsson & Jonsson, 2009) and that are currently co-occurring with the stressors investigated in this study, the consequences of these multiple interacting stressors on anadromous populations need further research. For this purpose, as we show in the present study, the description of the life history based on individual energetics is essential for a mechanistic understanding of their effects at the population and community levels. Our results show that in the face of multiple environmental threats the outcome of conservation efforts aimed at population persistence (i.e. increasing growth rate during oceanic life stage of Atlantic salmon) may in fact promote extinction and highlight the need for accurately predicting ecological consequences of environmental change. If we are to predict ecological consequences of environmental change a mechanistic understanding linking individual energy budgets, life history, and population dynamics will be almost certainly required.

## Supporting information

Model details, supporting information and figures

## Authors’ contribution

Chaparro-Pedraza analyzed the results and led the writing of the manuscript; de Roos contributed critically to the drafts. Both authors conceived the ideas, designed methodology, gave final approval for publication and agree to be held accountable for the work performed therein.

## Acknowledgements

The authors gratefully acknowledge the data courteously provided by the University of Idaho and the United States Army Corps of Engineers (SI Appendix, Fig. S2) and by B. Martin and R. B. MacFarlane (Fig. 1b), as well as P.C.de Ruiter for comments on the manuscript.

## Funding

This research was supported by the European Research Council (ERC) under the European Union’s Seventh Framework Programme (FP/2007-2013)/ ERC Grant Agreement no.322814.

## Supporting information appendix associated to this manuscript includes

Supporting information text, figures S1 to S5, table S1 and S2 and references for supporting information.

## References

Alexander, R. M. (1998). When is migration worthwhile for animals that walk, swim or fly? Journal of Avian Biology, 29(4), 387–394.

ASFB. (2010). Annual review. Retrieved from http://www.asfb.org.uk/wp-content/uploads/2011/04/ASFB-Review-2010.pdf

Auer, S. K., Arendt, J. D., Chandramouli, R., & Reznick, D. N. (2010). Juvenile compensatory growth has negative consequences for reproduction in Trinidadian guppies (Poecilia reticulata). Ecology Letters, 13(8), 998–1007. http://doi.org/10.1111/j.1461-0248.2010.01491.x

Bowerman, T. E., Pinson-Dumm, A., Peery, C. A., & Caudill, C. C. (2017). Reproductive energy expenditure and changes in body morphology for a population of Chinook salmon Oncorhynchus tshawytscha with a long distance migration. Journal of Fish Biology, 1–20. http://doi.org/10.1111/jfb.13274

Caudill, C. C., Daigle, W. R., Keefer, M. L., Boggs, C. T., Jepson, M. A., Burke, B. J., … Zabel, R. W. (2007). Slow dam passage in adult Columbia River salmonids associated with unsuccessful migration: delayed negative effects of passage obstacles or condition-dependent mortality? Can. J. Fish Aquat. Sci, 64, 979–995. http://doi.org/10.1139/F07-065

Crain, C. M., Kroeker, K., & Halpern, B. S. (2008). Interactive and cumulative effects of multiple human stressors in marine systems. Ecology Letters, 11(12), 1304–1315. http://doi.org/10.1111/j.1461-0248.2008.01253.x

Crossin, G. T., Hinch, S. G., Farrell, A. P., Higgs, D. A., Lotto, A. G., Oakes, J. D., & Healey, M. C. (2004). Energetics and morphology of sockeye salmon: Effects of upriver migratory distance and elevation. Journal of Fish Biology, 65(3), 788–810. http://doi.org/10.1111/j.1095-8649.2004.00486.x

de Roos, A. M. (1988). Numerical methods for structured population models: The Escalator Boxcar Train. Numerical Methods for Partial Differential Equations, 4(3), 173–195. http://doi.org/10.1002/num.1690040303

Eliason, E. J., Clark, T. D., Hague, M. J., Hanson, L. M., Gallagher, Z. S., Jeffries, K. M., … Farrell, A. P. (2011). Differences in thermal tolerance among Sockeye Salmon populations, (April), 109–113.

FAO. (2016). The state of world fisheries and aquaculture 2016. Contributing to food security and nutrition for all. Rome.

Friedland, K. D., MacLean, J. C., Hansen, L. P., Peyronnet, A. J., Karlsson, L., Reddin, D. G., … McCarthy, J. L. (2009). The recruitment of Atlantic salmon in Europe. ICES Journal of Marine Science, 66(2), 289–304. http://doi.org/10.1093/icesjms/fsn210

Gagliano, M., & McCormick, M. I. (2007). Compensating in the wild: Is flexible growth the key to early juvenile survival? Oikos, 116(1), 111–120. http://doi.org/10.1111/j.2006.0030-1299.15418.x

Glebe, B. D., & Leggett, W. C. (1981). Latitudinal differences in energy allocation and use during the freshwater migrations of American shad (Alosa sapidissima) and their life history consequences. Canadian Journal of Fisheries and Aquatic Sciences, 38(7), 806–820. http://doi.org/10.1139/f81-109

Hendry, K., & Cragg-Hine, D. (2003). Ecology of the Atlantic Salmon. Conserving Natura 2000 Rivers Ecology Series No. 7. English Nature. Peterborough.

Hoegh-Guldberg, O., & Bruno, J. F. (2010). The impact of climate change on the world’s marine ecosystem. Science, 328(June), 1523–1528. http://doi.org/10.1080/00330124.2015.1124788

ICOLD. (2017). World register of dams. Retrieved February 7, 2017, from www.icold-cigb.org

Johansen, S. J. S., Ekli, M., Stangnes, B., & Jobling, M. (2001). Weight gain and lipid deposition in Atlantic salmon, Salmo salar, during compensatory growth: Evidence for lipostatic regulation? Aquaculture Research, 32(12), 963–974. http://doi.org/10.1046/j.1365-2109.2001.00632.x

Jonsson, B., & Jonsson, N. (2004). Factors affecting marine production of Atlantic salmon (Salmo salar). Canadian Journal of Fisheries and Aquatic Sciences, 61(12), 2369–2383. http://doi.org/10.1139/f04-215

Jonsson, B., & Jonsson, N. (2009). A review of the likely effects of climate change on anadromous Atlantic salmon Salmo salar and brown trout Salmo trutta, with particular reference to water temperature and flow. Journal of Fish Biology, 75(10), 2381–2447. http://doi.org/10.1111/j.1095-8649.2009.02380.x

Jonsson, B., & Jonsson, N. (2016). Fecundity and water flow influence the dynamics of Atlantic salmon. Ecology of Freshwater Fish, (May), 1–6. http://doi.org/10.1111/eff.12294

Jonsson, N., Jonsson, B., & Hansen, L. P. (1997). Changes in proximate composition and estimates of energetic costs during upstream migration and spawning in Atlantic salmon Salmo salar. Journal of Animal Ecology, 66(3), 425–436. http://doi.org/10.2307/5987

Jonsson, N., Jonsson, B., & Hansen, L. P. (1998). The relative role of density-dependent and density-independent survival in the life cycle of atlantic salmon Salmo salar. Journal of Animal Ecology, 67(5), 751–762. http://doi.org/10.1046/j.1365-2656.1998.00237.x

Jutila, E., Jokikokko, E., & Julkunen, M. (2006). Long-term changes in the smolt size and age of Atlantic salmon, Salmo salar L., in a northern Baltic river related to parr density, growth opportunity and postsmolt survival. Ecology of Freshwater Fish, 15(3), 321–330. http://doi.org/10.1111/j.1600-0633.2006.00171.x

Kleinteich, A., Wilder, S. M., & Schneider, J. M. (2015). Contributions of juvenile and adult diet to the lifetime reproductive success and lifespan of a spider. Oikos, 124(2), 130–138. http://doi.org/10.1111/oik.01421

Kooijman, S. a. L. M. (2010). Dynamic Energy Budget theory for metabolic organisation (Vol. 1). http://doi.org/10.1098/rstb.2010.0167

Kooijman, S. A. L. M., & Metz, J. A. J. (1984). On the dynamics of chemically stressed populations: The deduction of population consequences from effects on individuals. Hydrobiological Bulletin, 17(1), 88–89. http://doi.org/10.1007/BF02255198

Lenders, H. J. R., Chamuleau, T. P. M., Hendriks, A. J., Lauwerier, R. C. G. M., Leuven, R. S. E. W., & Verberk, W. C. E. P. (2016). Historical rise of waterpower initiated the collapse of salmon stocks. Scientific Reports, 6, 1–9. http://doi.org/10.1038/srep29269

Limburg, K. E., & Waldman, J. R. (2009). Dramatic declines in North Atlantic diadromous fishes. BioScience, 59(11), 955–965. http://doi.org/10.1525/bio.2009.59.11.7

MacFarlane, R. B. (2010). Energy dynamics and growth of Chinook salmon (Oncorhynchus tshawytscha) from the Central Valley of California during the estuarine phase and first ocean year. Canadian Journal of Fisheries and Aquatic Sciences, 67(10), 1549–1565. http://doi.org/10.1139/F10-080

Martin, B., Heintz, R., Danner, E., & Nisbet, R. (2017). Integrating lipid storage into general representations of fish energetics. Journal of Animal Ecology, 86(2), 812–825. http://doi.org/10.1111/1365-2656.12667

Mesa, M., & Magie, C. (2006). Evaluation of energy expenditure in adult spring chinook salmon migrating upstream in the columbia river basin: an assessment based on sequential proximate analysis. River Research and Applications, 22, 1085–1095. http://doi.org/10.1002/rra

Moore, J. K., Fu, W., Primeau, F., Britten, G. L., Lindsay, K., Long, M., … Randerson, J. T. (2018). Sustained climate warming drives declining marine biological productivity. Science, 359(6380), 1139–1143. http://doi.org/10.1126/science.aao6379

Persson, L., Leonardsson, K., de Roos, a M., Gyllenberg, M., & Christensen, B. (1998). Ontogenetic scaling of foraging rates and the dynamics of a size-structured consumer-resource model. Theoretical Population Biology, 54(3), 270–293. http://doi.org/10.1006/tpbi.1998.1380

Sinervo, B., & Doughty, P. (1996). Interactive effects of offspring size and timing of reproduction on offspring reproduction: Experimental, maternal, and quantitative genetic aspects, 50(3), 1314–1327.

Taborsky, B. (2006). The influence of juvenile and adult environments on life-history trajectories. Proceedings. Biological Sciences / The Royal Society, 273(1587), 741–750. http://doi.org/10.1098/rspb.2005.3347

Thorpe, J. E., Miles, M. S., & Keay, D. S. (1984). Developmental rate, fecundity and egg size in Atlantic Salmon, Salmo Salar L. Aquaculture, 43, 289–305. http://doi.org/10.1016/0044-8486(84)90030-9

Walters, A. W., Copeland, T., & Venditti, D. A. (2013). The density dilemma: Limitations on juvenile production in threatened salmon populations. Ecology of Freshwater Fish, 22(4), 508–519. http://doi.org/10.1111/eff.12046

WCD. (2000). Dams and Development: A new framework for decision-making. Earthscan Publications.

Werner, E. E., & Gilliam, J. F. (1984). The Ontogenetic Niche and Species Interactions in Size-Structured Populations. Ecology, 15, 393–425.

Zeller, M., & Koella, J. C. (2016). Effects of food variability on growth and reproduction of Aedes aegypti. Ecology and Evolution, 6(2), 552–559. http://doi.org/10.1002/ece3.1888

