## Supplementary material for "Low marine food levels mitigate high migration costs in anadromous populations": Model details, supporting information and figures

This supplementary note presents a detailed population-level model based on the individual life history described in the methods section of the main text. It also presents evidence and theory supporting two essential assumptions of the model: 1) the habitat switch to higher food levels results in larger and leaner individuals, 2) large individuals spend disproportionally more energy than small ones transporting their body upstream. Likewise, we show the effect of a nonlinear scaling of the costs of the breeding migration with respect to structural mass and of including the reversible mass in the costs of the breeding migration.

*Physiologically–structured population model*

The physiologically structured population model follows the approach introduced by Persson et al.(1) for populations with seasonal reproduction, in which the population is represented by a dynamic set of cohorts or year classes. Since reproduction occurs as a discrete event at a specific time in the year, all individuals that are born in the same reproductive event are considered equal and hence lumped into a single cohort and assumed to grow at the same rate. Thus, we can describe the dynamics of each cohort $i \in \mathbb{N}$ by using a system of ordinary differential equations, which keeps track of the density of individuals $N_i$, their age $A_i$, their structural mass $W_i$ and their reversible mass $S_i$. Juveniles are defined as individuals with structural mass smaller than the structural mass at maturity $W_p$ and adults as individuals with structural mass equal or larger than $W_p$. For each cohort $i$, age is monotonically increasing with time,

$$\frac{d}{dt}A_i = 1 \tag{1}$$

The age of the individuals determines the stage, which in turn, determines the differential equations that describe the variation in density of individuals, their structural mass and stored energy reserves. Equations (2), (3) and (4) below define the dynamics of eggs, presmolts and postsmolts respectively. The number of individuals decreases due to a mortality rate specific to each stage. In addition, the presmolts and postsmolts may die due to starvation. During the egg stage the structural mass and storage does not change. The dynamics of the structural and reversible mass in presmolts and postsmolts depend on the amount of food they encounter as well as the breeding migration period if they are adults.

for $0 \leq A_i < a_h$

$$\begin{cases} \frac{d}{dt} N_i = -\mu_e N_i \\ \frac{d}{dt} W_i = 0 \\ \frac{d}{dt} S_i = 0 \end{cases}$$

(2)

for $a_h \leq A_i < a_s$

$$\begin{cases} \frac{d}{dt} N_i = \begin{cases} -\mu_r N_i & if\ \frac{S_i}{W_i} \geq q_S \\ -\left(\mu_r N_i + \varphi\left(q_S \frac{W_i}{S_i} - 1\right)\right) & if\ S_i > 0\ and\ \frac{S_i}{W_i} < q_S \\ -\infty & otherwise \end{cases} \\ \frac{d}{dt} W_i = \begin{cases} \zeta_W \left(\kappa\ \frac{R_r}{K + R_r}\ j_a\ {W_i}^{2/3} - j_m\ W_i\right) & if\ \kappa\ \frac{R_r}{K + R_r}\ j_a\ {W_i}^{2/3} > j_m\ W_i \\ 0 & otherwise \end{cases} \\ \frac{d}{dt} S_i = \begin{cases} (1 - \kappa) \frac{R_r}{K + R_r}\ j_a\ {W_i}^{2/3} & if\ \kappa\ \frac{R_r}{K + R_r}\ j_a\ {W_i}^{2/3} > j_m\ W_i \\ \frac{R_r}{K + R_r} j_a\ {W_i}^{2/3} - j_m\ W_i & otherwise \end{cases} \end{cases}$$

for $a_s \leq A_i$

$$\begin{cases} \frac{d}{dt}N_i = \begin{cases} -\mu_s N_i & if\ \frac{S_i}{W_i} \geq q_S \\ -\left(\mu_s N_i + \varphi\left(q_S \frac{W_i}{S_i} - 1\right)\right) & if\ S_i > 0\ and\ \frac{S_i}{W_i} < q_S \\ -\infty & otherwise \end{cases} \\ \frac{d}{dt}W_i = \begin{cases} \zeta_W\left(\kappa\ f_s\, j_a\ W_i^{2/3} - j_m\ W_i\right) & if\ c1\ and\ (\sim c2\ or\ \sim c3) \\ 0 & otherwise \end{cases} \\ \frac{d}{dt}S_i = \begin{cases} (1-\kappa) f_s\, j_a\ W_i^{2/3} & if\ c1\ and\ (\sim c2\ or\ \sim c3) \\ f_s\, j_a\ W_i^{2/3} - j_m\, W_i & if\ \sim c1\ and\ (\sim c2\ or\ \sim c3) \\ -(j_m\ W_i + C\, j'_m\ W_i^{PW}) & otherwise \end{cases} \end{cases}$$

(4)

In this last equation $c1$, $c2$ and $c3$ represent the conditions $\kappa\ f_s\, j_a\ W_i^{2/3} > j_m\ W_i$, $t_{um} \leq t \leq t_{dm}$, and $W_p \leq W_i$ respectively, while $\sim c1$, $\sim c2$ and $\sim c3$ refer to the situation that these conditions do not hold. When the conditions are true, the $\kappa$ fraction of the amount of assimilates necessary is sufficient to meet metabolic maintenance ($c1$), the current time corresponds to the breeding migration period ($c2$) and the cohort is adult ($c3$).

Whenever a juvenile cohort reaches the maturation size $W_i = W_p$, at a particular time $t = t_p$, a maturation event occurs. At a maturation event, the juvenile cohort becomes an adult cohort. This does not affect any cohort statistics:

$$\begin{cases} A_i(t_p) = A_i(t_p^-) \\ N_i(t_p) = N_i(t_p^-) \\ W_i(t_p) = W_i(t_p^-) \\ S_i(t_p) = S_i(t_p^-) \end{cases}$$

(5)

Reproduction occurs instantaneously at $t = n\ t_y + t_r$, where $n \in \mathbb{N}$. At a reproductive event, a new cohort is formed from the reversible biomass of adults, if their reversible:structural mass ratio exceeds the reversible:structural mass ratio with which the adults matured:

$$\begin{cases} A_0(t_{rn}) = 0 \\ N_0(t_{rn}) = \left( \sum_{i \in \{j \le n | W_j \ge W_p\}} N_i \cdot \max\left( S_i - \frac{S_p}{W_p} W_i, 0 \right) \right) \frac{\zeta_e}{W_e} \\ W_o(t_{rn}) = \kappa\, W_b \\ S_0(t_{rn}) = (1-\kappa)\, W_b \end{cases}$$

(6)

At the same time, all other cohorts are renumbered and the reversible mass of the reproducing adults is set to the amount that makes their reversible:structural mass ratio equal to their reversible:structural mass ratio at maturation.

$$\begin{cases} A_{i+1}(t_{rn}) = A_i(t_{rn}{}^{-}) \\ N_{i+1}(t_{rn}) = N_i(t_{rn}{}^{-}) \\ W_{i+1}(t_{rn}) = W_i(t_{rn}{}^{-}) \\ S_{i+1}(t_{rn}) = \begin{cases} \min\left( S_i(t_{rn}{}^{-}), \frac{S_p}{W_p} W_i(t_{rn}{}^{-}) \right) & if\ W_i \ge W_p \\ S_i(t_{rn}{}^{-}) & otherwise \end{cases} \end{cases}$$

(7)

The resource density in the breeding habitat grows following a semi–chemostat growth and declines by foraging of presmolts (8).

$$\frac{d}{dt} R_r = \rho(R_{max} - R_r) - \frac{R_r}{K + R_r} j_a \sum_{i \in \{j \le n | a_h < a_j < a_s\}} N_i\, W_i^{2/3}$$

(8)

*Energy allocation effects of the habitat switch explained by dynamic energy budget theory*

Dynamic energy budget (DEB) theory provides a conceptual framework to describe the individual life history based on individual energetic dynamics. DEB theory describes the rules by which an individual assimilates energy and utilizes it to grow, reproduce and cover metabolic maintenance (2–4). It has been used to describe the life history of several species including salmonids (5). In particular, the net assimilation model offers a conceptual explanation for the negative effect on fecundity caused by an increase in food abundance. This model assumes that a fraction $\kappa$ of assimilates is allocated to first meet metabolic maintenance with the remainder of this fraction allocated to growth in structural mass, while a fraction $1 - \kappa$ is allocated to growth in reversible mass to be used for reproduction (Fig S1a) and covering energetic deficits during starvation periods. Given the assumption that metabolic maintenance is deducted from the fraction $\kappa$, a change in the proportion of assimilates required to meet metabolic maintenance affects the proportion of assimilates allocated to growth in structural mass but not the fraction allocated to growth in reversible mass. When an individual experiences a step-up change in food, the amount of assimilates available for growth, reproduction and metabolic maintenance increases. However, the amount of assimilates required to meet metabolic maintenance remains constant because the somatic structure of the individual does not suddenly change. Since the amount of total assimilates increases, the proportion of assimilates to meet metabolic maintenance thus decreases with the surplus now being allocated to growth in structural mass. Therefore, the proportion of assimilates allocated to growth in structural mass increases, while the proportion of assimilates allocated to growth in reversible mass, and thus to reproduction remains constant (Fig S1b). The model hence predicts that an individual that experiences a step-up change in food has lower energy density (lower ratio of reversible to structural mass) and consequently, lower mass-specific fecundity than an individual of the same size that never experiences a change in food, in line with data presented in Fig 1. Furthermore, the model predicts that this bias toward increased growth in structural mass compared to reversible mass is larger in individuals experiencing a large step-up change in food than in those experiencing a small one. Consequently, individuals that experience a large change in food grow larger (are bigger) and have a lower energy density (are leaner).

*Size-scaling of the breeding migration costs with structural mass and breeding migration costs dependent on structural and reversible mass*

Metabolic rates of swimming and resting salmon scale with body mass with a power 0.79 and 0.78, respectively (6). It is important to notice that these exponents hold for an allometric relation between metabolism and body mass, which quantities are different from the metabolic maintenance and structural mass in the model that correspond to only part of the metabolism and part of the body mass, respectively. However, this study suggests that metabolic costs of swimming scale with a similar factor than resting metabolic costs. Nonetheless, this study is based on fish swimming in still water but for swimming against a current the energetic costs are different because the optimal speed is higher in the latter than in the former (6).

The breeding migration entails multiple costs. In the freshwater habitat where the habitat is not iso-osmotic, individuals invest energy in osmoregulation, such that respiration rate can increase more than 20% just due to osmoregulatory expenses (7). These osmoregulatory costs increase in proportion to the surface area of individual (3) and therefore scale allometrically with the structural mass $W$ with an exponent equal to 0.67. In addition, large fishes travel upstream using portions of the river further from the bank than small ones (8) where the current is faster and therefore they spend more energy traveling against a faster current (9). In support of these arguments, data of energy expenditure during the breeding migration of Chinook salmon *Oncorhynchus tshawytscha* show that the size-specific energy requirements of large individuals are larger than of small individuals (10). That is, larger individuals spend more energy per unit of structural mass than smaller ones (Fig S2), hence $PW$ is larger than 1. The energy loss during the migratory travel is also higher in large than in small individuals of Atlantic salmon (11) and American Shad (*Alosa sapidissima*) (12).

Given this, we evaluated the effect that different size-scaling exponents of the costs of the breeding migration $PW$ have on population persistence (Fig S3). Smaller values of $PW$ increase persistence when costs of the breeding migration are high, because reversible mass is depleted to a lesser extent during the breeding migration. Persistence of a migratory population at low food levels but extinction occurring for higher food levels in the ocean when the costs of the breeding migration are high occurs when $PW$ is 0.5 or larger. This effect is reversed for smaller PW, for example when equal to 0.3. However, the evidence presented above makes values of $PW$ below 0.5 unlikely and suggests that it actually scales with a value larger than 1 with respect to body size, resulting in a stronger persistence effect of declining food in the ocean when a population faces high costs of the breeding migration because large individuals have higher size-specific migration costs than small individuals. Based on that evidence, a choice of the size-scaling exponent of the energetic costs of the breeding migration $PW$ equal to 1 with respect to the structural mass as used in Fig 2 and 3 is conservative. Our assumption of a size-scaling exponent of the energetic costs of the breeding migration $PW$ of 1 implies that structural mass-specific energetic costs of the breeding migration are the same for every individual regardless of their body size, while energetic costs of the breeding migration per unit of total mass (structural plus reversible) decrease with body size because the reversible mass increases with body size (Fig S4).

An increasing impact of reversible mass on the costs of breeding migration, meaning taking into account the total body mass in these costs, has no qualitative effects for the increase of persistence of a migratory population at low food levels but extinction occurring for higher food levels in the ocean when the costs of the breeding migration are high. However, the costs of the breeding migration at which this phenomenon occurs are lower compared to the case in which the costs of the breeding migration are independent of the reversible mass. When the costs of the breeding migration are proportional to structural mass only the population shows a small variation in the population biomass when the decrease of food level in the ocean occurs in year 20, whereas when the costs of the breeding migration are proportional to total mass (structural plus reversible) the population increases in biomass in response to the decrease of food level (Fig S5). Therefore, including the reversible mass in the costs of the breeding migration causes the population experiencing high food levels in the ocean to go extinct at lower costs of the breeding migration but still to persist at low food levels in the ocean.

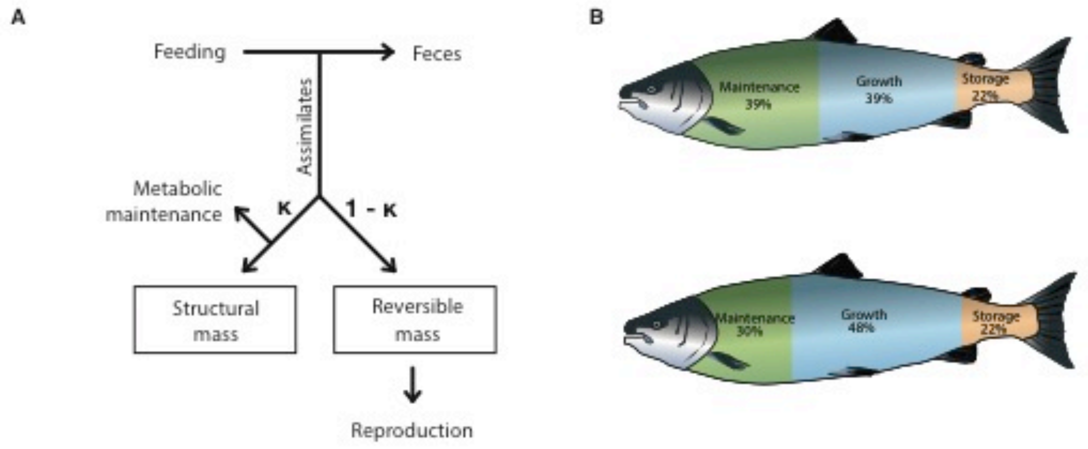


**Figure S1**. Energy allocation rule causes bias toward somatic growth after a large step-up change in food

**A:** Net assimilation energy budget model. When a step-up change in food occurs, metabolic maintenance requirements remain constant because the somatic structure only changes after growth has occurred. Because a fixed fraction $\kappa$ of assimilates is allocated to cover both metabolic maintenance requirements and growth in structural mass, the sudden increase in available assimilates translates into a proportionally larger increase in the allocation to growth in structural mass compared to the increase in reversible mass. **B:** Proportion of assimilates allocated to growth (in structural mass), storage (reversible mass) and metabolic maintenance of individuals exposed to either a small (feeding level changes from 50 to 60% of maximum food intake; top) or large step-up in food abundance when switching habitats (feeding level changes from 50 to 90% of maximum food intake; bottom). See also Fig 2 and energy allocation effects of the habitat switch explained by dynamic energy budget theory.

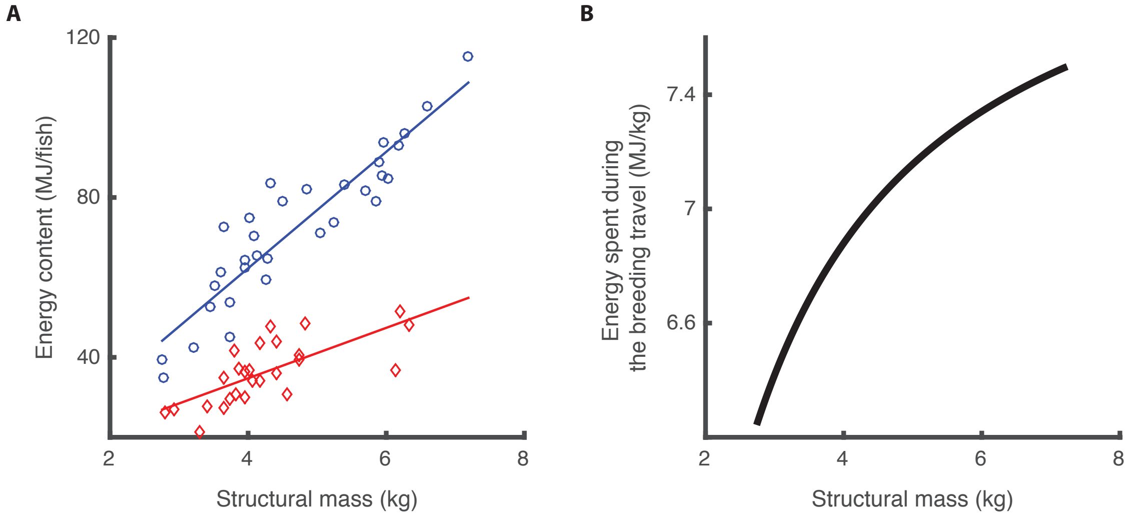


**Figure S2**. Title: Nonlinear scaling of the costs of the breeding travel with size in wild sockeye salmon

**A:** Total energy content of female individuals of *Onchorhynchus tshawytscha* at the beginning of the breeding travel (Ebt = 14.57 Structural mass + 4.0085, R-squared = 0.845, blue) and at arrival in the spawning grounds (Eat = 6.3057 Structural mass + 9.5457, R-squared = 0.514, red) from Bowerman et al(10) (data courteously provided by T. Bowerman). Structural mass was calculated from fork length Lf data ($\text{Structural mass} = \text{dw}\ (\text{Lf} * \text{sc})^3$, where $\text{dw} = 1\ \text{g cm}^{-3}$ is the density of the organism and $\text{sc} = 0.2$ is the shape coefficient for this species (5). **B:** Mass-specific energy expenditure calculated as the difference between the total energy content at the beginning of the breeding travel and at arrival in spawning grounds, and divided by the structural mass ($\text{Mass\_specific energy spent during the breeding travel} = 8.26 - 5.54\ (\text{somatic mass}^{-1})$). See also Size-scaling of the breeding migration costs with structural mass and breeding migration costs dependent on structural and reversible mass.

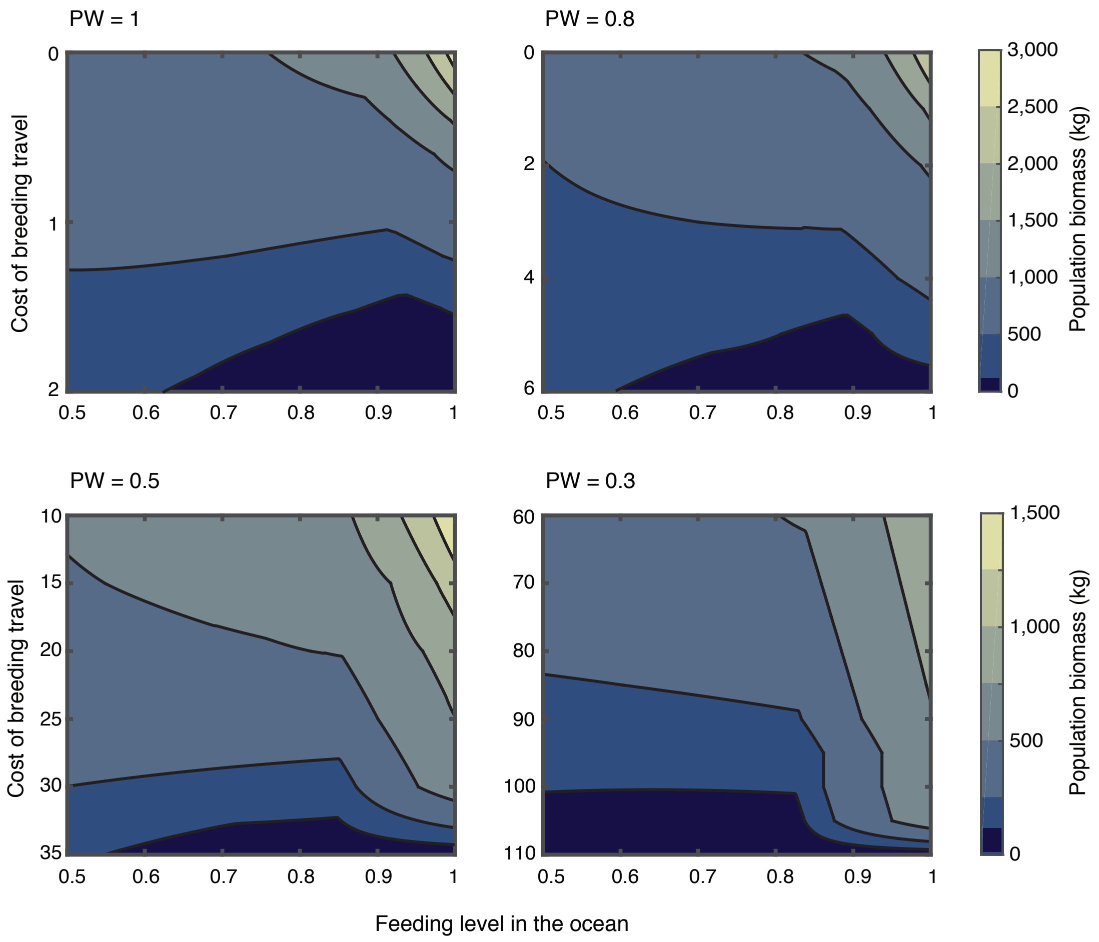


**Figure S3**. Consequences of interacting threats for population biomass with nonlinear scaling of the costs of the breeding travel with body size

Predicted biomass of anadromous populations exposed to different feeding levels in the ocean (horizontal axes) and facing different costs of the breeding travel (vertical axes) for four different scaling exponents of the costs of the breeding travel with individual body size. A population is considered to be extinct when its biomass is smaller than 100 kg (dark blue). Values shown represent the average population biomass computed over the stable annual cycle that the populations exhibit. See also Size-scaling of the breeding migration costs with structural mass and breeding migration costs dependent on structural and reversible mass.

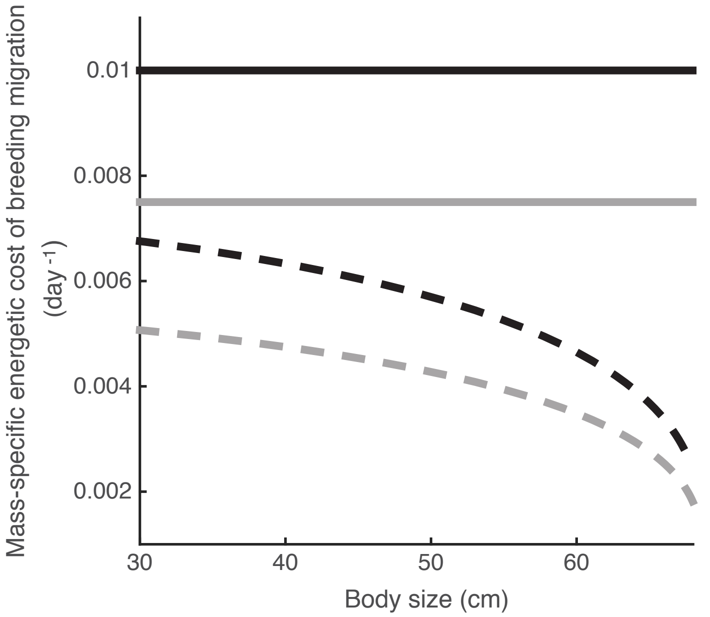


**Figure S4**. Mass-specific costs of the breeding migration when this energetic costs scales with a factor of 1 with respect to structural mass ($PW = 1$) and the relative costs of the breeding migration are 0.5 ($C = 0.5$, grey lines) and 1 ($C = 1$, black lines).

For this factor, the energetic cost of the breeding migration divided by structural mass is the same for every individual regardless of its body size (solid lines), while the energetic cost of the breeding migration divided by its total body mass (structural mass + reversible mass) decreases with body size because reversible mass increases with body size. Data are representative for an individual migrating to the breeding grounds for first time after a feeding level of 0.6 in the ocean ($f_s = 0.6$).

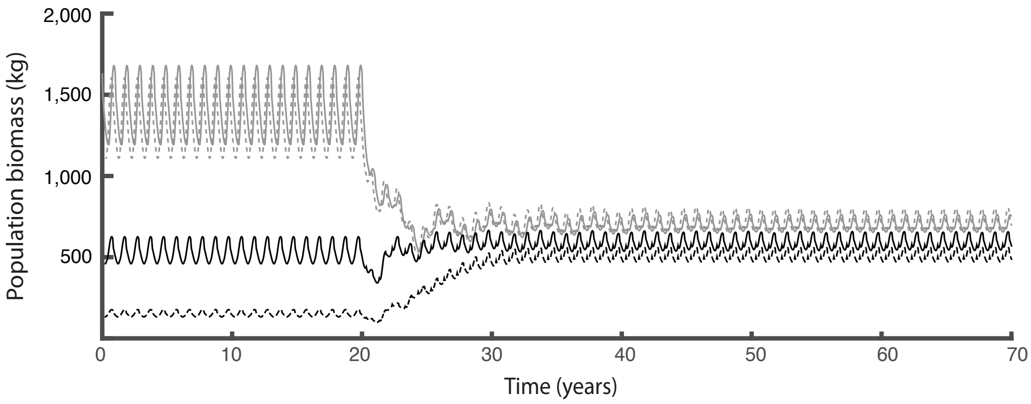


**Figure S5**. Population consequences of declining food abundance in the ocean when migration costs are dependent on total mass (structural plus reversible)

Biomass dynamics of a population facing low (0.5 times the normal field metabolic costs; grey) and high (1 times the normal field metabolic costs; black) costs of the breeding travel preceding and following a drop in feeding level in the ocean (as in Fig 2). Solid lines show dynamics for the default case (eq. 12) and dotted lines show the dynamics when the costs of the breeding travel dependent on total (structural plus reversible) mass (eq. 15). Notice that in the latter case the increase of population biomass when decreasing food levels in the ocean occurs at lower energy costs than in the former (Fig 2). See also Size-scaling of the breeding migration costs with structural mass and breeding migration costs dependent on structural and reversible mass.

**Table S1.** Summary of results of compensatory growth studies that reported fecundity

| Species | Growth | Reduced fecundity due to step-up change? | Average individual fecundity | | Units | Reference |
|---|---|---|---|---|---|---|
| | | | After step-up change in food | Control | | |
| *Daphnia magna* (Cladoceran) | Indeterminate | Yes | 15.1 * | 60.1 * | eggs | (Kooijman, unpublished) |
| *Poecilia reticulata* (Fish) | Indeterminate | Yes | 40.6 * | 52.3 * | eggs | (13) |
| *Phalloptycus januarius* (Fish) | Indeterminate | Yes | 4.5 * | 7.5 * | eggs/week | (14) |
| *Uta stansburiana* (Lizard) | Indeterminate | Yes | 3.53 * | 5.1 * | eggs/ clutch | (15) |
| *Aedes aegypti* (Insect) | Determinate | Yes | 49 * | 70 * | eggs | (16) |
| *Larinioides sclopetarius* (Arachnid) | Determinate | Yes | 384 ** | 772 ** | eggs | (17) |
| *Coturnix coturnix* (Bird) | Determinate | No | | | | (18) |
| *Mus musculus* (Mammal) | Determinate | No | | | | (19) |

*Data digitalized from figures in the original publication

**Data listed in the original publication

**Table S2.** Parameter values

| Description | Symbol | Value | Unit | References |
|---|---|---|---|---|
| Environment | | | | |
| Year | $t_y$ | 365 | day | |
| Average temperature | $T_m$ | 283 | K | |
| Amplitude of temperature variation | $T_a$ | 278 | K | |
| Events within the season | | | | |
| Day of the beginning of breeding travel | $t_{um}$ | 205 | | (20) |
| Day of reproduction (spawning) | $t_r$ | 215 | | (21) |
| Day of the end of breeding travel | $t_{dm}$ | 225 | | (20) |
| Age-dependent events during life cycle | | | | |
| Age at hatching | $a_h$ | 150 | day | (21,22) |
| Age at smolting | $a_s$ | 545 | day | (23) |
| Food resource in the breeding habitat | | | | |
| Resource growth rate | $\rho$ | 0.1 | $day^{-1}$ | |
| Resource maximum density | $R_{max}$ | 5 | $g\ m^{-3}$ | |
| Half saturation resource density | $K$ | 1 | $g\ m^{-3}$ | |
| Migratory population | | | | |
| Feeding level of postsmolts | $f_s$ | varied | - | |
| Fraction of assimilation flux to structural mass growth and maintenance | $\kappa$ | 0.8 | | (5,24) |
| Maximum area-specific assimilation rate | $j_a$ | 0.18 | $g\ g^{-2/3}\ day^{-1}$ | Calculated with method of Jager (25) from regressions of Koskela et al (26) |
| Mass-specific metabolic maintenance costs | $j_m$ | 0.006 | $g\ g^{-1}\ day^{-1}$ | Calculated with method of Jager (25) from regressions of Koskela et al (26) |
| Mass-specific metabolic costs of the breeding travel | $j'_m$ | 0.006 | $g\ g^{-PW}\ day^{-1}$ | |
| Reference temperature | $T^*$ | 293 | K | |
| Arrhenius temperature | $T_A$ | 8000 | K | |
| Yield of structural mass on assimilates | $\zeta_W$ | 0.8 | $g\ g^{-1}$ | (25) |
| Yield of egg buffer on reversible mass | $\zeta_e$ | 0.95 | $g\ g^{-1}$ | (25) |
| Mass of a single egg | $W_e$ | 0.1 | g | (27) |
| Mass of a new born (after hatching) | $W_b$ | 0.06 | g | (28) |
| Structural mass at maturity | $W_p$ | 74 | g | (5) |
| Shape coefficient factor | $\delta$ | 0.21 | - | (5) |
| Density of structural mass | $v$ | 1 | $g\ cm^{-3}$ | |
| Costs of the breeding travel | $C$ | varied | - | |
| Size scaling exponent of the costs of the breeding travel | $PW$ | varied | - | |
| Mortality rate of eggs | $\mu_e$ | 0.0125 | $day^{-1}$ | (29) |
| Mortality rate of presmolts | $\mu_r$ | 0.0025 | $day^{-1}$ | (30) |
| Mortality rate of postsmolts | $\mu_s$ | varied | $day^{-1}$ | |
| Minimum reversible/structural mass ratio that individuals stand without starvation mortality | $q_S$ | 0.1 | - | (1) |
| Scaling coefficient for starvation mortality | $\varphi$ | 0.2 | - | (1) |

### References


1. Persson L, Leonardsson K, de Roos a M, Gyllenberg M, Christensen B. Ontogenetic scaling of foraging rates and the dynamics of a size-structured consumer-resource model. Theoretical population biology. 1998;54(3):270–93.

2. Kooijman SALM, Metz JAJ. On the dynamics of chemically stressed populations: The deduction of population consequences from effects on individuals. Hydrobiological Bulletin. 1984;17(1):88–9.

3. Kooijman S a. LM. Dynamic Energy Budget theory for metabolic organisation [Internet]. 2010. 514 p. Available from: http://www.pubmedcentral.nih.gov/articlerender.fcgi?artid=2981979&tool=pmcentrez&rendertype=abstract

4. Nisbet RM, Muller EB, Lika K, Kooijman SALM. From molecules to ecosystems through dynamic energy budget models. Journal of Animal Ecology. 2000;69(6):913–26.

5. Pecquerie L, Johnson LR, Kooijman SALM, Nisbet RM. Analyzing variations in life-history traits of Pacific salmon in the context of Dynamic Energy Budget (DEB) theory. Journal of Sea Research. 2011;66(4):424–33.

6. Alexander RM. When is migration worthwhile for animals that walk, swim or fly? Journal of Avian Biology. 1998;29(4):387–94.

7. Fry FEJ. The effect of environmental factors on the physiology of fish. Fish Physiology. 1971;6:1–98.

8. Hughes NF. The wave-drag hypothesis: an explanation for size-based lateral segregation during the upstream migration of salmonids. Canadian Journal of Fisheries and Aquatic Sciences. 2004;61(1):103–9.

9. Hinch SG, Rand PS. Optimal swimming speeds and forward-assisted propulsion: energy-conserving behaviours of upriver-migrating adult salmon. Can J Fish Aquat Sci. 2000;57:2470–8.

10. Bowerman TE, Pinson-Dumm A, Peery CA, Caudill CC. Reproductive energy expenditure and changes in body morphology for a population of Chinook salmon *Oncorhynchus tshawytscha* with a long distance migration. Journal of Fish Biology [Internet]. 2017;1–20. Available from: http://doi.wiley.com/10.1111/jfb.13274

11. Jonsson N, Jonsson B, Hansen LP. Changes in proximate composition and estimates of energetic costs during upstream migration and spawning in Atlantic salmon Salmo salar. Journal of Animal Ecology. 1997;66(3):425–36.

12. Glebe BD, Leggett WC. Latitudinal differences in energy allocation and use during the freshwater migrations of American shad (Alosa sapidissima) and their life history consequences. Canadian Journal of Fisheries and Aquatic Sciences. 1981;38(7):806–

20.

13. Auer SK, Arendt JD, Chandramouli R, Reznick DN. Juvenile compensatory growth has negative consequences for reproduction in Trinidadian guppies (Poecilia reticulata). Ecology Letters. 2010;13(8):998–1007.

14. Pollux BJA, Reznick DN. Matrotrophy limits a female's ability to adaptively adjust offspring size and fecundity in fluctuating environments. Functional Ecology. 2011;25(4):747–56.

15. Sinervo B, Doughty P. Interactive Effects of Offspring Size and Timing of Reproduction on Offspring Reproduction : Experimental , Maternal , and Quantitative Genetic Aspects. 1996;50(3):1314–27.

16. Zeller M, Koella JC. Effects of food variability on growth and reproduction of Aedes aegypti. Ecology and Evolution. 2016;6(2):552–9.

17. Kleinteich A, Wilder SM, Schneider JM. Contributions of juvenile and adult diet to the lifetime reproductive success and lifespan of a spider. Oikos. 2015;124(2):130–8.

18. Hassan SM, Mady ME, Cartwright AL, Sabri HM, Mobarak MS. Effects of early feed restriction on some performance and reproductive parameters in Japanese Quail ( Coturnix coturnix japonica ). International Journal of Poultry Science 13. 2003;13(6):323–8.

19. Zamiri MJ. Effects of reduced food intake on reproduction in mice. Australian Journal of Biological Sciences. 1978;31:629–39.

20. Doucett RR, Booth RK, Power G, McKinley RS. Effects of the spawning migration on the nutritional status of anadromous Atlantic salmon (*Salmo salar*): insights from stable-isotope analysis. Canadian Journal of Fisheries and Aquatic Sciences. 1999;56(11):2172–80.

21. Thorpe JE, Mangel M, Metcalfe NB, Huntingford F a. Modelling the proximate basis of salmonid life-history variation, with application to Atlantic salmon, Salmo salar L. Evolutionary Ecology. 1998;12(5):581–99.

22. Hendry K, Cragg-Hine D. Ecology of the Atlantic Salmon. Conserving Natura 2000 Rivers Ecology Series No. 7. English Nature. Peterborough; 2003.

23. Mccormick SD, Hansen LP, Quinn TP, Saunders RL. Movement, migration, and smolting of Atlantic salmon (Salmo salar). Canadian Journal of Fisheries and Aquatic Sciences. 1998;55:77–92.

24. Jager T, Martin BT, Zimmer EI. DEBkiss or the quest for the simplest generic model of animal life history. Journal of Theoretical Biology [Internet]. Elsevier; 2013;328:9–18. Available from: http://dx.doi.org/10.1016/j.jtbi.2013.03.011

25. Jager T. DEBkiss A simple framework for animal energy budgets. 2015;

26. Koskela J, Pirhonen J, Jobling M. Feed intake, growth rate and body composition of juvenile Baltic salmon exposed to different constant temperatures. Aquaculture

international. 1997;5:351–60.

27. Potts WTW, Rudy PP. Water balance in the eggs of the Atlantic salmon Salmo salar. The Journal of Experimental Biology. 1969;(50):223–37.

28. Shearer K, Asgard T, Andorsdottir G, Aas G. Whole body elemental and proximate composition of Atlantic salmon (Salmo salar) during the life cycle. Journal of Fish Biology. 1994;44:785–97.

29. Bley PW, Moring JR. Freshwater and Ocean Survival of Atlantic Salmon and Steelhead: A Synopsis [Internet]. 1988. Available from: http://oai.dtic.mil/oai/oai?verb=getRecord&metadataPrefix=html&identifier=ADA322801

30. Leggett WC, Power G. Differences Between Two Populations of Landlocked Atlantic Salmon (Salmo salar) in Newfoundland. Journal of the Fisheries Research Board of Canada [Internet]. NRC Research Press; 1969 Jun 1;26(6):1585–96. Available from: http://dx.doi.org/10.1139/f69-142
